# Environmental enrichment in piglet creeps: behavior and productive performance

**DOI:** 10.1101/346023

**Authors:** Karina Sartor, Bolivar Felipe de Freitas, Juliana de Souza Granja Barros, Luiz Antonio Rossi

**Affiliations:** University of Campinas – UNICAMP – School of Agricultural Engineering, Av. Cândido Rondon 501, 13083875 Campinas, Brazil

**Keywords:** Environmental enrichment, creeps, piglets, animal welfare, sensory stimuli

## Abstract

In the farrowing stage, to use heated creeps is crucial for meeting the thermal requirements of newborn piglets and alternatives for environmental enrichment to attract the animals to the creeps. The objective of this study was to evaluate the behavior and productive performance of piglets submitted to creeps enriched with different types of sensory stimuli. The study had a completely randomized design. The animals were submitted to seven treatments: **T1**) Enrichment with blue LED lighting; **T2**) Enrichment with fragrance of chamomile (*Matricaria chamomilla*) essential oil; **T3**) Enrichment with fragrance of lavender (*Lavandula*) essential oil; **T4**) Enrichment with fragrance of lemon (*Citrus* × *latifolia*) essential oil; **T5**) Enrichment with classical music, “The Four Seasons” by Vivaldi; **T6**) Creeps heated without environmental enrichment (control group); **T7**) Enrichment with fragrance of thyme (*Thymus vulgaris*) essential oil. The piglets were submitted to creeps with environmental enrichment under automatic control every 15 min (on/off), from 8 am to 5:45 pm (21 consecutive days). The environmental conditions of the creeps – air temperature, relative humidity, luminous intensity, and decibels – were evaluated within the creeps with environmental enrichment to determine the influence on the piglets’ behavior and productive performance. The results show that piglets submitted to creeps with environmental enrichment with blue artificial light and thyme essential oil showed reduced frequency outside the creeps when compared with treatment T6 (control). This showed the reduction of the stay of piglets in areas susceptible to crushing, in proximities to the mother. Piglets showed greater preference for creeps enriched with fragrance diffusion of thyme essential oil and with blue artificial lighting.

## Introduction

In the intensive rearing of pigs, one of the major problems is to meet the thermal requirements and attract the piglets into heated creeps using environmental enrichment to avoid hypothermia, low weight gain, and high mortality rate.

The most critical period for survival of piglets is up to 72 hours after birth, resulting from the following variables: birth weight, birth order, air temperature, behavior, deaths by crushing, starvation, and exposure to cold [1]. Heated creeps can attract the piglets away from areas with danger of crushing close to the mother, reducing the likelihood of crushing [2]. Since newborn piglets, by instinct, seek to lay along with the mother to maintain thermoregulation, they are very prone to death by crushing due to their small size when compared with the mother [3].

The use of heated creeps meets the needs of protection and comfort. However, the training for animals to identify this environment is still performed manually: piglets must be placed inside the creep one by one, and must be locked up a few times a day, in the first days of life. Newborn piglets need to be trained, in the first 24 hours of life, to access the creeps several times [4]. In this process, it is important to determine whether the piglets are able to make an active choice based on their individual preferences. Through the training, the piglets learn to use the creeps from the 3rd day of age after birth [5,6].

To prevent piglets from seeking thermal comfort with their mother, the use of environmental enrichment of creeps through sensory stimuli (aromatization, lighting, and music) can arouse curiosity and exploration of the thermal environment comfortable to them.

Newborn piglets are attracted by thermal, olfactory, visual, and tactile stimuli [7,8]. Fragrances can be one of these resources, showing favorable characteristics both due to the ability of arousing the piglet’s attraction to the environment [9,10] and to its biological effects and effect of sanitization of the environment, and thus may contribute to improve the health of the animals. Examples of fragrances that can be used and that have these favorable characteristics include essential oils extracted from plants [11].

Visual stimulation of piglets using artificial lighting can influence their investigative behavior. [12] observed that young piglets prefer exploring illuminated environments, rather than those without lighting. [13] demonstrated that piglets exposed to music in the farrowing stage had decreased aggressive behavior after weaning. [14] observed higher consumption and weaning weight of pigs submitted to musical sound. [11] emphasized that the use of music has relaxing effect on humans and animals.

Thus, it is necessary to seek alternatives for environmental enrichment through sensory stimuli that arouse the piglets’ attraction to the heated creeps in conditions of thermal comfort.

This study was developed with the objective of evaluating the behavior and productive performance of piglets submitted to creeps enriched with different types of sensory stimuli.

## Material and methods

The experiment was conducted in Holambra, SP, Brazil, at latitude 22°37’59”S and longitude 47°03’20”W and altitude of 600 meters over sea level. The climate of the region is humid subtropical (Cfa) according to the Köppen classification.

The research followed the ethical standards and was approved by the Ethics Committee on Animal Use (registration No. 4317-1) of the University of Campinas, SP.

The study used 18 farrows of the Landrace breed (230 piglets), all from the same genetic lineage at the farrowing stage. The piglets were submitted to creeps (1 m long by 0.56 m wide and 0.54 m high, polyethylene material, 5 mm thick) heated (heating established by the farm management – 100 W lamp) with and without environmental enrichment, in the period of 21 days (January 4 to 25, 2017).

Visual (blue lighting), olfactory (fragrance of essential oils), and auditory (music) sensory stimuli were chosen based on results documented in the literature for the welfare of humans and other animals, since there are few scientific studies on the effect of sensory stimuli on pigs.

The swine identified the blue artificial lighting (visual stimuli) (440–490 nm) [15]; the aromatization of the environment with infusion of volatile essential oils (olfactory stimulation) of lavender (decreases anxiety), Tahiti lime (sedative effect in humans) [16], Chamomile (soothing effect for dogs) [17], and thyme (antibacterial effect [18] and muscle relaxing effect for mice [19]); and classical music (auditory stimuli) [20] that has soothing effect [21], and to which swine respond at frequency of up to 85 dB [14].

The sows (n=21) were distributed into 7 treatments in completely randomized design with about 32 piglets in each treatment (mean of 10.6 piglets/repetition). The treatments were:

**Treatment – T1:** Enrichment with blue LED lighting;

**Treatment – T2:** Enrichment with fragrance of chamomile (*Matricaria chamomilla*) essential oil;

**Treatment – T3:** Enrichment with fragrance of lavender (*Lavandula*) essential oil;

**Treatment – T4:** Enrichment with fragrance of lemon (*Citrus* × *latifolia*);

**Treatment – T5:** Enrichment with classical music, “The Four Seasons” by Vivaldi;

**Treatment – T6:** Heated creeps without environmental enrichment (control group).

**Treatment – T7:** Enriched with fragrance of thyme (*Thymus vulgaris*) essential oil.

The creeps were installed with speakers (speaker, micro SD slot, and auxiliary jack – WS-887), electric diffuser (DWY125V-S) adapted for diffusion of essential oil, and blue LED Lamp (SMD3528). A time switch (RTST/20) was used for the automatic programming of environmental enrichment (blue led lighting and electric diffuser) every 15 min (on/off) in the creeps, from 8 am to 5:45 pm.

Environmental enrichment of the creeps with classical music, The Four Seasons, composed by Antonio Vivaldi, was programmed by means of the Zara Radio software with the aid of the computer. The time switch and the Zara Radio software were programmed to trigger the environmental enrichment of the creeps in the same time interval of the other treatments. Figure 1 (A, B, and C) shows the installation of environmental enrichment within the creeps:

**Fig 1.**
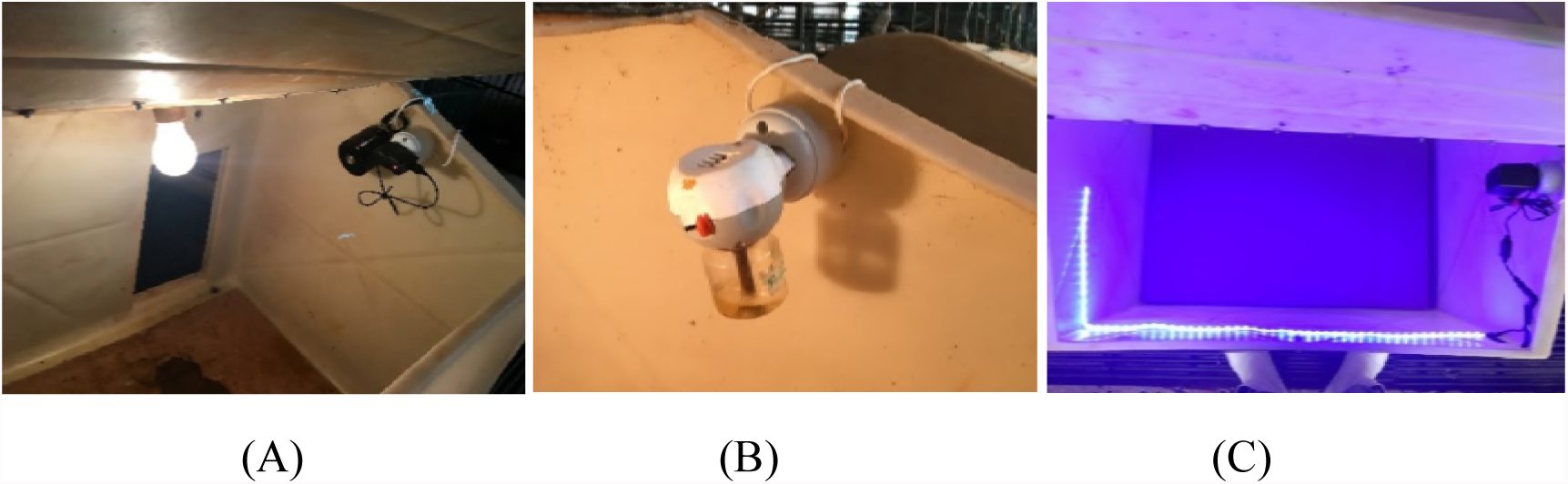
Speaker (A), essential oil fragrance diffuser (B), LED lighting (C).

Data on air temperature (°C), relative humidity (%), luminous intensity (lx), and sound pressure (dB) of creeps with environmental enrichment were evaluated to determine the influence on piglet’s behavior (%) and piglet’s performance (kg).Temperature and relative humidity (USB Datalogger, model TM-305U accuracy ±0.6°C and ±3% RH) were registered within the creeps every 10 minutes. Luminous intensity (digital lux meter LUX METER, Model *Lx*-101) and noise level in decibels (model ITDEC-3000, accuracy±2 dB) were measured on 3 random days at 9 am and 3 pm.

Piglets’ behavior was evaluated based on exposure to the environmental enrichment of the creeps. Piglets’ behaviors were measured through direct observation [13], in the morning at 8 am, 9 am, and 10 am and in the afternoon at 2 pm, 3 pm, and 4 pm, in the period of 10 random days (total of 1,080 observations). Piglets’ behaviors in the adaptation period after birth (first 3 days of life) were not considered. Therefore, the piglets’ behavioral responses of thermal comfort, exploratory profile, and agonistic profile were measured in their activities. Table 1 shows the ethogram of behavioral observations of piglets.

**Table 1.** . Behavioral ethogram used to evaluate the piglets’ thermal comfort, exploratory profile, and agonistic profile.

| Behavior | Description |
| --- | --- |
| Lying scattered | Piglets sleeping, lying far from each other on the heating source, indicating thermal comfort. |
| Lying on top of each other | Piglets sleeping lying in groups when they are cold to exchange heat and maintain homeothermy. |
| Suckling | The piglet's act of suckling the sow. |
| Lying beside the mother | Piglets lying beside the mother to maintain homeothermy. |
| Frequency of piglets | Piglets within and outside the creep. |
| Agonistic profile | Fights, struggles, and bites between the piglets. |
| Exploratory profile | Piglet's interaction with environment (smelling, nudging another piglet or components in the creep and pen). |

[Adapted from 22, 5, 23]

The influence of environmental enrichment in creeps on piglet productive performance was evaluated through the initial weight (based on the end of transfer management of piglets according to the number of functional teats of sows) and weight at weaning (final weight). The number of piglet deaths was measured in each treatment for calculation of mortality rate.

The means of the evaluated variables were submitted to analysis of variance (ANOVA) followed by Tukey test (p<0.05) with the aid of statistical software Statgraphics Centurion XV.

## Results

### Environmental variables

In the farrowing room, the mean air temperature value at 24.19°C and relative humidity at 90% during the experimental period (21 days) was maintained through evaporative cooling system. The values presented in Table 2 compare the means for luminous intensity in lux (lx), sound pressure in decibels (dB), air temperature (°C), and relative humidity (%) between the treatments.

**Table 2.** Values for environmental variables on luminous intensity (lx), sound pressure (dB), air temperature (°C), and relative humidity (%).

| Environmental variables |  |  |  |  |
| --- | --- | --- | --- | --- |
| Treatments | lx | dB | °C | RH |
| T1 | 803.15 ($\pm 124.49$)<sup>a</sup> | 69.45 ($\pm 1.44$)<sup>b</sup> | 29.70 ($\pm 0.42$)<sup>a</sup> | 62.28 ($\pm 2.93$)<sup>a</sup> |
| T2 | 689.63 ($\pm 103.41$)<sup>b</sup> | 67.65 ($\pm 2.55$)<sup>b</sup> | 29.70 ($\pm 0.29$)<sup>a</sup> | 61.57 ($\pm 1.41$)<sup>a</sup> |
| T3 | 673.52 ($\pm 117.71$)<sup>b</sup> | 67.97 ($\pm 3.36$)<sup>b</sup> | 29.64 ($\pm 0.66$)<sup>a</sup> | 62.43 ($\pm 4.28$)<sup>a</sup> |
| T4 | 643.25 ($\pm 79.33$)<sup>b</sup> | 66.83 ($\pm 2.78$)<sup>b</sup> | 29.20 ($\pm 1.05$)<sup>a</sup> | 63.13 ($\pm 0.86$)<sup>a</sup> |
| T5 | 686.63 ($\pm 116.28$)<sup>b</sup> | 77.12 ($\pm 4.09$)<sup>a</sup> | 30.25 ($\pm 0.25$)<sup>a</sup> | 61.61 ($\pm 1.13$)<sup>a</sup> |
| T6 | 578.00 ($\pm 169.54$)<sup>b</sup> | 68.49 ($\pm 2.45$)<sup>b</sup> | 29.02 ($\pm 0.31$)<sup>a</sup> | 64.83 ($\pm 0.54$)<sup>a</sup> |
| T7 | 654.20 ($\pm 117.26$)<sup>b</sup> | 68.07 ($\pm 2.51$)<sup>b</sup> | 29.58 ($\pm 0.49$)<sup>a</sup> | 62.64 ($\pm 1.85$)<sup>a</sup> |

Same letters, in the column, do not differ statistically by Tukey test ( $p>0.05$ ). Mean ± standard deviation.
**lx**=luminous intensity; **dB**=decibels; **RH**=relative humidity. Treatments: **T1** – blue lighting; **T2** – Chamomile essential oil diffusion; **T3** – Lavender essential oil diffusion; **T4** – Lemon essential oil diffusion; **T5** – classical music; **T6** – Control; **T7** – Thyme essential oil diffusion.

Luminous intensity (T1) differed (p<0.05) between treatments T2, T3, T4, T5, T6, and T7. In the group with blue lighting treatment (T1), the lux was of 803.15 lx, value higher than treatments T2 (689.63 lx), T3 (673.52 lx), T4 (643.25 lx), T5 (686.63 lx), T6 (578.00 lx), and T7 (654.20 lx). Sound pressure (dB) in treatment T5 (music) differed (p<0.05) between the treatments (T1, T2, T3, T4, T6, and T7). Air temperature and relative humidity did not differ (p>0.05) between treatments because of the same thermal condition provided to animals in creeps with and without environmental enrichment (Table 2). Figure 2 presents the mean values for piglets’ behavior frequency inside creeps, in each treatment.

**Fig 2.**
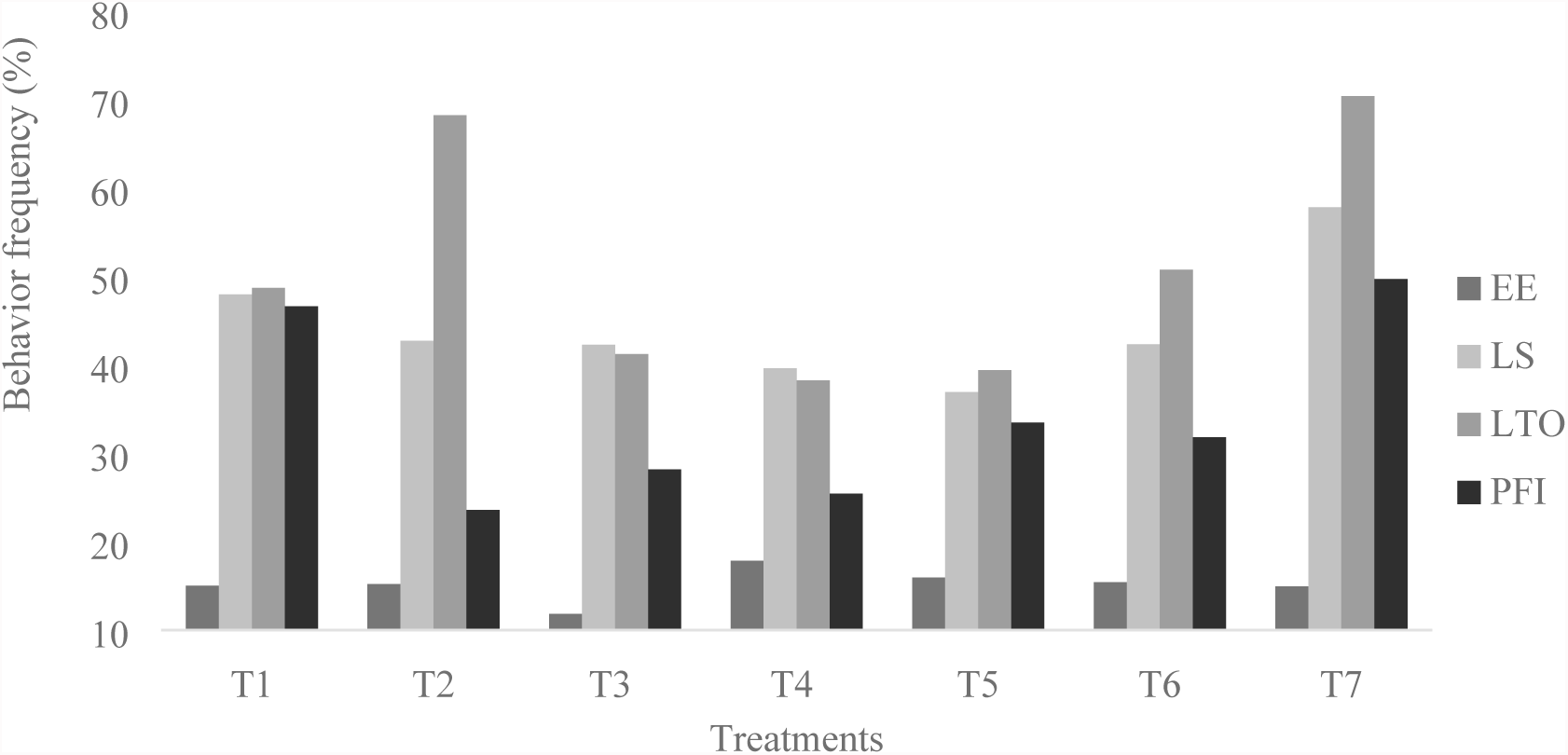
Mean values (%) for piglet behavior frequency in creeps with and without environmental enrichment. Behaviors: EE – exploring the creep; LS – lying scattered; LTO – lying on top of each other; PFI - piglets within the creep. Treatments: T1 – blue Lighting; T2 – Chamomile essential oil diffusion; T3 – Lavender essential oil diffusion; T4 – Lemon essential oil diffusion; T5 – classical music; T6 – Control; T7 – thyme essential oil diffusion.

The piglets’ frequency of exploratory behavior and behavior of lying on top of one another inside the creep did not differ statistically (p>0.05) between any of the treatments (Fig. 2), and none of the respective treatments influenced the behaviors of the piglets. The agonistic behavior of the piglets inside the creep was limiting for this phase because the animals explore a comfortable area inside the creep to lie down and rest.

The behavioral frequency of piglets lying scattered inside the creep differed statistically (p<0.05) in treatment T7 (aromatization with fragrance of Thyme essential oil), compared with treatments T2, T3, T4, T5, and T6. The frequency of behaviors of lying scattered differed (p<0.05) between treatments T1 (blue lighting) and T5 (classical music), showing that the piglets became active when submitted to musical stimuli. The piglets showed higher frequency for the behavior of lying scattered in treatment T7 (57.84%) in creep enriched with fragrance of Thyme essential oil, compared with the other treatments. Treatments T1 and T7 did not differ statistically (p>0.05) compared with each other and showed similar mean values for behavior of piglets lying scattered, validating the piglets’ behavioral response of welfare in both treatments. The lower behavioral frequency of piglets lying scattered was 36.94% for group T5, indicating that the piglets remained more active and less rested when submitted to stimulation of classical music. To rest, piglets preferred the environment with artificial blue lighting over the sound of classical music. The total behavioral frequency of piglet’s access into the creeps differed (p<0.05) in treatment T7 compared with treatments T2, T3, T4, T5, and T6 (Table 2). Piglet’s access in treatment T1 differed (p<0.05) compared with treatments T6, T5, T4, and T2. However, piglets submitted to environmental enrichment with Thyme essential oil (T7) and artificial blue lighting (T1) showed better effect compared with the control treatment T6 (without environmental enrichment).

Figure 3 shows the behaviors: exploratory profile, agonistic profile, suckling, lying beside mother, and total access of animals outside the creeps.

**Fig 3.**
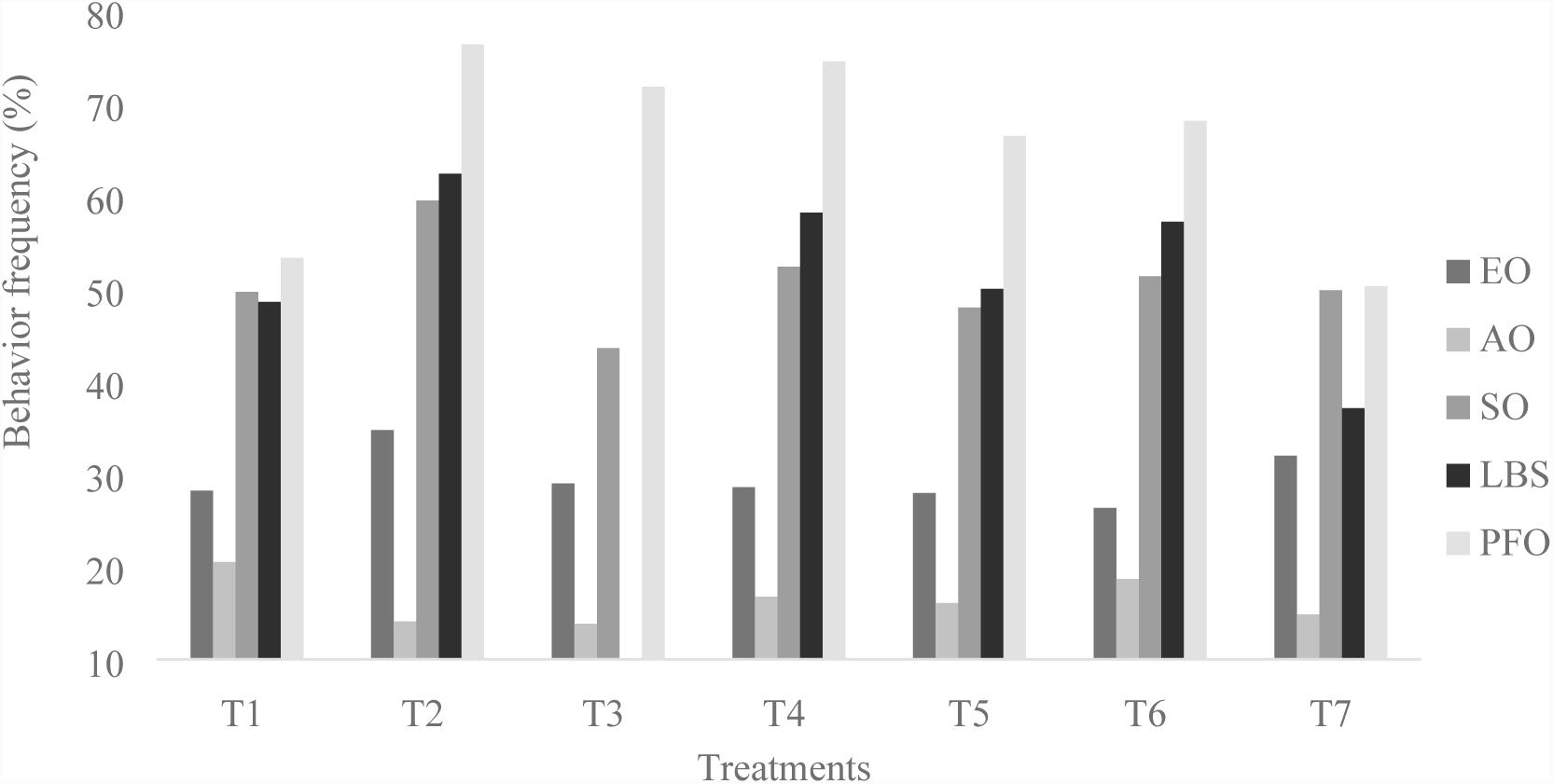
Mean values for piglets’ behavior frequency outside the creep. Behaviors: EO – exploratory profile outside the creep; AO – agonistic profile outside creep; SO – suckling; LBS – lying beside the sow; PFO – piglets outside creep. Treatments: T1 – blue LED Lighting; T2 – Chamomile essential oil diffusion; T3 – Lavender essential oil diffusion; T4 – Lemon essential oil diffusion; T5 – classical music; T6 – control; T7 – Thyme essential oil diffusion.

The frequency behaviors of exploratory profile, agonistic profile, and lying beside the mother outside the creeps, in the pen area, did not differ statistically (p>0.05) between all treatments. The frequency of piglet suckling differed (p<0.05) in treatment T2 (Chamomile) compared with treatments T3 (Lavender) and T6 (control group). Piglets showed higher frequency for suckling in the treatment enriched with Chamomile essential oil Chamomile essential oil (59.56%) and Lavender (43.63%) compared with the control treatment (51.37%) without use of environmental enrichment (Fig 3). However, the increased frequency of piglet suckling did not result in better piglet performance (Fig. 3). The total frequency of piglets (Fig. 3) outside the creeps differed (p<0.05) between treatments T4 (Lemon essential oil diffusion) and T7 (Thyme essential oil diffusion). The frequency of the piglets’ behavior outside the creep did not differ (p>0.05) between treatments T2, T3, T4, T5, and T6, showing that the environmental enrichments used (diffusion with essential oil of Chamomile, Lavender, Lemon, and classical music) in the creeps were not attractive for piglets compared with control treatment (T6).

Piglets submitted to treatments T1 and T7, enriched with artificial blue lighting (53.27%) and Thyme essential oil (50.28%), showed lower frequency outside the creeps compared with treatment T6 (control). These results (Fig 3) show the reduction of the stay of piglets in areas susceptible to crushing, close to the mother, with use of artificial blue lighting and Thyme essential oil diffusion in creeps.

### Piglet’s productive performance

Weaning weight of piglets submitted to the treatments was related to the thermal conditions of heating to which the animals were subjected. Table 3 shows the weaning weight of piglets submitted to the treatments.

**Table 3.**
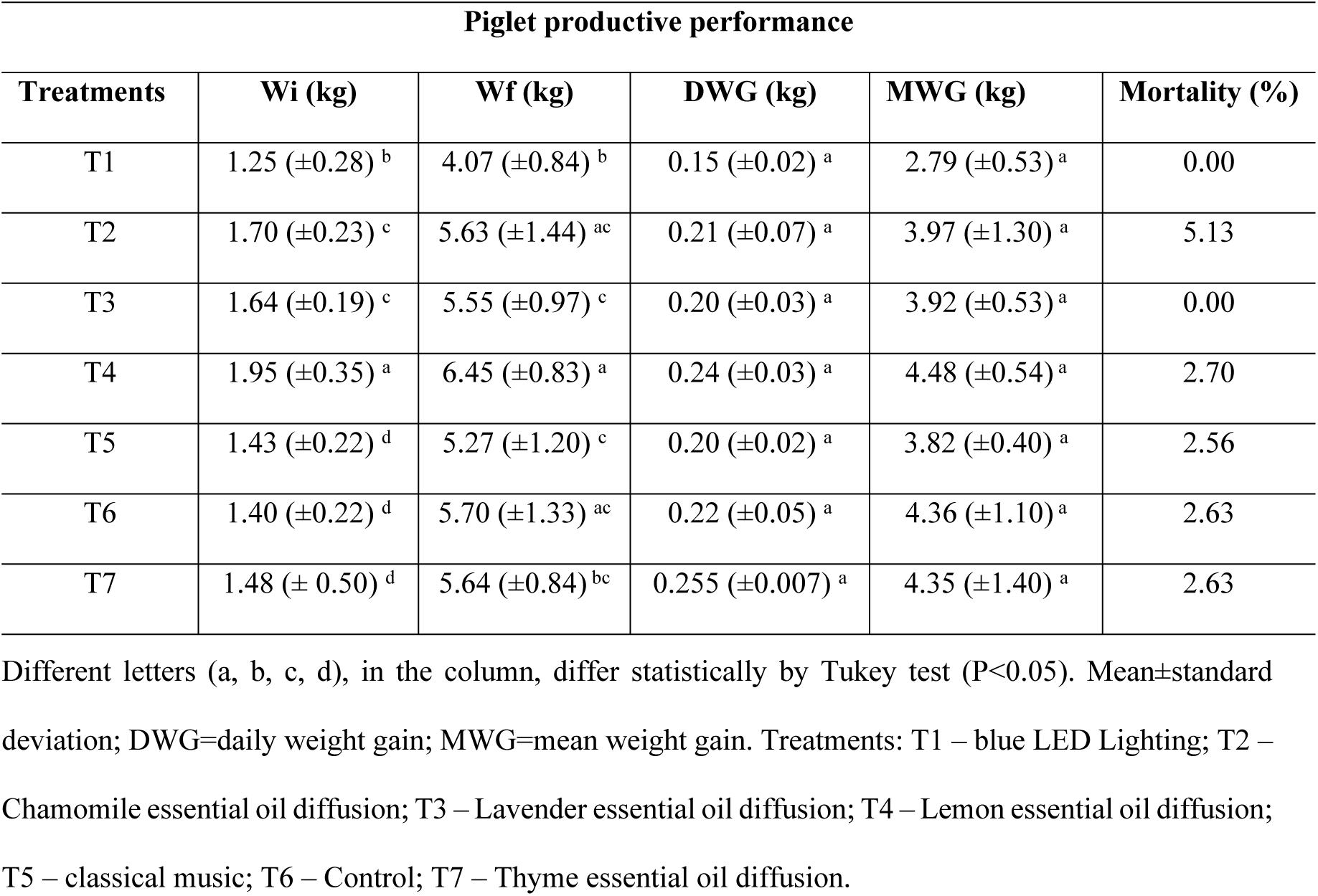
Mean values for initial weaning weight (Wi), final body weight (Wf), daily weight gain (DWG), mean weight gain (MWG) of the piglets submitted to the treatments.

The piglets’ initial body weight in treatment T4 (1.95 kg) differed statistically (p<0.05) compared with treatments T1 (1.25 kg), T2 (1.70 kg), T3 (1.64 kg), T5 (1.43 kg), T6 (1.40 kg), and T7 (1.48 kg). Differently, the final body weight at weaning differed (p<0.05) in treatment T4 (6.45 kg), compared with treatments T1 (4.07 kg), T3 (5.55 kg), T5 (5.27 kg), and T7 (5.64 kg). The piglets’ daily weight gain (DWG) and mean weight gain (MWG) in the period did not differ (p>0.05) in any of the treatments.

## Discussion

In treatment T1, the increase in the mean value for lux (803.00 lx) was influenced by the incandescent lamp (100 W) used as heating and by the artificial blue lighting (Table 2). Luminous intensity in treatments T2, T3, T4, T5, T6, and T7 was maintained between 578.00 and 686.63 lx, influenced by the incandescent lamp. There are few scientific studies concerning the amount of lighting indicated for pigs. [24] indicated 80 lux to ensure animal welfare, while the European legislation [25] recommends a minimum of 40 lux. According to [26], providing additional lighting for swine influences on their behavior, making them more active in their activities. [27,28] claim that animals react to the contrast of light and tend to opt for a more illuminated area. [12] observed that young piglets prefer illuminated environments over dark ones, as they appear to be afraid of the latter.

The increase in the mean value of decibels in treatment T5 is due to the environmental enrichment with classical music (Table 2). The noise level was below the recommended for swine, 84 dB, and brought no risk to the welfare of piglets [29]. Piglets submitted to environmental enrichment with musical sound at a frequency level between 80 and 85 dB showed better performance compared with the environment without music [14].

Experiments were conducted (groups submitted to different types of environmental enrichment and control group) in creeps with the same thermal condition (heating management established by the farm), close to the minimum air temperature of thermal comfort (30°C) for piglets (Table 2). Piglets must be maintained in the air temperature range between 30 and 34°C to avoid hypothermia [4] and obtain satisfactory weaning gain. Relative humidity between the treatments remained in the range of thermal comfort between 50 and 80%, adequate thermal conditions and relative humidity for swine [30]. Creep environmental enrichment with fragrance diffusion of thyme essential oil increased the behavioral frequency of piglets lying scattered and the frequency of access into the creeps (Fig. 2), characteristics that indicate attraction to the environmental enrichment. Studies show that piglets show behavioral episodes in lying scattered position in comfortable environments [22,31]. Piglets also lie scattered or on top of one another to sleep and rest.

Piglets submitted to environmental enrichment with classical music showed lower frequency in behavior (Fig. 2) of rest (lying scattered). However, we observed that the noise level and the rhythm or sound frequency can cause scare reactions in piglets [14] or stimulate playful behavior [13], making them more active.

The frequencies of behaviors of exploration and of lying on top of each other inside the creeps were correlated between the treatments, because the air temperature remained homogenous in both treatments (Fig. 2). Studies prove that piglets seek to lie down, quickly, in groups when the air temperature decreases below the thermal comfort range [32]. The piglets’ exploratory behavior was consistent with that observed in the study of [33], in which they spent about 10–15% of the time exploring the environment. This shows that the animals in confinement show limited natural behavior, even with the use of environmental enrichment.

Piglets showed preference for accessing creeps enriched with artificial blue lighting (T1 – 46.63%) and with the fragrance of Thyme essential oil (T7 – 49.72%), compared with the other experimental groups (Fig. 2). Even with luminous intensity above 80 lx in all treatments, the piglets showed greater preference for creeps with artificial blue lighting (T1 – 803.15 lx) than in treatments T2, T3, T4, T5, and T6. These results suggest that piglets can identify the blue lighting. [34] support the theory that the swine have rods and cones in the retina with sensitive structures for detection of wavelengths of blue and green colors in the visible spectrum. For [35], the swine identify and distinguish blue (440–490 nm) from the other colors but cannot perceive the colors red and green. [36] claim that the wavelengths identified by swine species range from 439 to 556 nm, classified as the color blue (440-490 nm). [37] tested colors in water troughs and found that swine were attracted by red or blue objects and ignored the green object.

Piglets submitted to fragrance diffusion of Thyme essential oil were more attracted (49.72%) in entering the creeps compared with the other essential oils used as environmental enrichment and with the control treatment without environmental enrichment. This result could be due to the composition and concentration of the chemical constituents volatilized by diffusion of essential oil fragrances absorbed through the respiratory tract of piglets. Chemical constituents released by diffusion of essential oils can act on neurotransmitter activation, producing the feeling of calmness and welfare for animals in creeps.

In swine rearing, there are few scientific studies on the use of essential oils regarding the welfare effect of animals. A study with humans by [38] showed that inhalation of a small dose of essential oil activates the olfactory system by the olfactory nerves and bulb, with action on the Central Nervous System and activates brain regions responsible for social and behavioral emotions, and production of neurotransmitters, such as serotonin, acetylcholine, norepinephrine, endorphins and others that make communication with all systems of the body.

Environmental enrichment by diffusing fragrance of essential oil of Chamomile, Lavender, and Lemon had no effect on the frequency of piglet’s access to the creep (Fig. 2) compared with control treatment without environmental enrichment (T6). The smell of essential oils of lavender, lemon and chamomile may have caused a repulsive effect associated with the toxicity of foods or fear of predators, since essential oils can release, by diffusion, multiple chemical constituents, which may be unattractive to piglets.

The wild swine used the sense of smell instinctively to find food, detect potential predators or prey, and mark territory. Thus, smells related to food of wild swine were presented as lure for hunting, including animal and vegetable oils, considered attractive [39]. [40] observed that the use of straw bed with lavender smell during transport of pigs decreased the malaise, without reducing stress, measured by salivary cortisol concentrations.

On the other hand, studies show that the lack of attractiveness for piglets to stay within the enriched creeps can be related with the loss of interest in the kind of environmental enrichment, which limits its usefulness [41]. [33] concluded that to provide environmental enrichment in alternating periods of the day to increase piglet’s interest is ineffective, but to stimulate them through rewards can increase their curiosity.

[42] claims that the swine are motivated to express and to perform behaviors that are rewarding to them. [43] report that the valuation of the swine by environmental enrichment will depend on several factors, including the properties of the material itself to allow the investigation and manipulation and the degree of novelty. [3] observed that the most attractive environmental enrichment to pigs are ingestible, flavored, and chewable objects, since these animals are curious and show investigative behavior in search of food.

The piglets’ agonistic and exploratory behaviors outside the creeps referred to investigating, rooting, smelling, and interacting between siblings, which had no effect as to the different types of environmental enrichment in the creeps.

Piglets submitted to environmental enrichment showed higher frequency in suckling behaviors and in total piglets outside the creeps in treatment T2, indicating that the use of Chamomile essential oil was unattractive for piglets to attend the creep (Fig. 3). The piglets preferred to remain less frequently outside the creep, close to the mother, in the creeps enriched with Thyme essential oil and artificial blue lighting. This result was satisfactory to reduce the stay of piglets in areas with risk of crushing, close to the mother. However, it is known that the stay of piglets close to their mothers leads to increase in mortality caused by crushing. [44] concluded that piglets maintained in environments without environmental enrichment showed higher frequency in the behavior of rooting and chewing directed to components of the pen, manipulation of the udder of the mother and of the siblings. In another study, [24] emphasize that piglets lie close to the mother to establish a bond, seek warmth, stay in comfort, feed, and protect themselves.

In this experiment, piglets were weaned at 21 days of age and showed better final body weight at weaning in treatment T4 (6.45 kg) in comparison between treatments T1 (4.07 kg), T3 (5.55 kg), T5 (5.27 kg), and T7 (5.64 kg). Piglets submitted to treatment T4 had the best initial body weight (1.95 kg) and thus resulted in the greatest final weight gain at weaning in treatment T4 (6.45 kg). The results for the final mean weight at weaning corroborate the results found by [45], who evaluated the effect of piglets’ weaning age and found that piglets weaned at 21 days of age weighed about 6.10 kg. [46] evaluated the body weight of piglets weaned at early age (weaning before 18 days) and moderate age (weaning between 18 and 27 days) and emphasized that piglets weaned at early age weighed 5.20 kg and at moderate age weighed 6.07 kg at weaning. The values found (Table 3) for the daily weight gain of piglets in treatments T2, T3, T4, T5, T6, and T7 are similar to those found by [47] in which piglets obtained mean daily weight gain of 0.221 kg in the period of lactation.

Mortality rate was higher (5.13%) in treatment T2 using Chamomile essential oil in creep environmental enrichment, resulting from the increased frequency of suckling behavior (Fig. 2), which had an increase in mortality rate due to the proximity of the piglets to the mother when suckling. According to [23], the cause of piglets’ mortality by crushing by the sow is 6.4%. Thus, inadequate management and discomfort in creep environment lead piglets to lie down close to sows, which predisposes to crushing [48]. Another factor that predisposes to crushing of piglets is low birth weight [49]. This study enabled analyzing the positive effect for piglets submitted to treatment T1 (artificial blue lighting), which showed lower birth weight and resulted in 0% of deaths. For piglets submitted to the other environmental enrichments in creeps, mortality rate was similar (2.56 and 2.70%) between the treatments (T4, T5, T6, and T7).

## Conclusion

The effect of environmental enrichment in creeps using sensory stimuli to arouse the curiosity of piglets was proven. The piglets showed greater preference for creeps enriched with fragrance diffusion Thyme essential oil and with blue artificial lighting. Thus, olfactory (Thyme essential oil diffusion) and visual (blue lighting) stimuli reduce the stay of piglets in areas susceptible to crushing, in the proximities of the sow.

## Acknowledgments

The authors are thankful to FAPESP, CAPES and FAEPEX-UNICAMP for the financial support.

